# Significant temporal shifts on clonal and plasmid backgrounds of *Enterobacteriaceae* producing acquired AmpC in Portuguese clinical settings

**DOI:** 10.1101/553164

**Authors:** T. G. Ribeiro, Â. Novais, C. Rodrigues, R. Nascimento, F. Freitas, E. Machado, L. Peixe

## Abstract

**OBJECTIVES:** To provide detailed molecular data on clinical acquired AmpC (qAmpC)-producing *Enterobacteriaceae* from two different periods (2002-2008 and 2010-2013) in order to clarify the contribution of clonal and plasmid genetic platforms for the current epidemiological scenario concerning extended-spectrum beta-lactams resistance.

**METHODS:** We analysed 1246 *Enterobacteriaceae* non-susceptible to third-generation cephalosporins from 2 hospitals and 1 community laboratory between 2010 and 2013. Bacterial identification, antibiotic susceptibility, identification of qAmpC and plasmid-mediated quinolone resistance genes, clonal (PFGE, MLST) and plasmid (S1-/I-*Ceu*I-PFGE, replicon typing, hybridization) analysis were performed by standard methods. WGS was performed in two ST11-*K. pneumoniae* isolates harbouring DHA-1.

**RESULTS:** The occurrence of qAmpC was lower (2.6%) than that observed in a previous survey (7.4%), and varied slightly over time. Isolates produced DHA-1 (53%), CMY-2 (44%) or DHA-6 (3%), but significant epidemiological changes were observed in the two surveys. While DHA-1 persisted in different institutions by selection of a worldwide epidemic IncR plasmid in a ST11 harbouring KL105, CMY-2 rates increased over time linked to IncI1 plasmids (instead of IncK or IncA/C_2_) in multiple *E. coli* clones.

**CONCLUSIONS:** The higher frequency of DHA-1 qAmpC in these species contrasts with the scenario of most European countries. Furthermore, the different genetic backgrounds associated with either ESBL or qAmpC in our country might have contributed to their differential expansion.

## 1. Introduction

The worldwide contemporary epidemiology of *Enterobacteriaceae* resistant to broad-spectrum cephalosporins seems to be largely dominated by extended-spectrum β-lactamases (ESBL) than acquired AmpC β-lactamases (qAmpC) **[1]**. However, the occurrence and clinical impact of *Enterobacteriaceae* producing qAmpC is probably underestimated, since qAmpC producers are difficult to identify by conventional methods and qAmpC expression is frequently masked by other resistance mechanisms **[2, 3]**. Recent studies described an increase of specific qAmpC enzymes among clinical *Enterobacteriaceae* **[1, 2, 4]**, mostly CMY-2-producing *Escherichia coli* associated either with hospital or community infections and DHA-1-producing *Klebsiella pneumoniae* especially linked to nosocomial outbreaks **[2, 3, 5, 6]**.

In Portugal, relevant epidemiological changes regarding resistance to extended-spectrum β-lactams in *Enterobacteriaceae* have been observed in the last decades, including: i) escalating resistance rates to third-generation cephalosporins, ii) long-term dissemination of qAmpC (mainly in *K. pneumoniae*), iii) emergence and dispersal of plasmid-mediated carbapenemase resistance and iv) the shift from *E. coli* to *K. pneumoniae* as the main species associated with this problem (https://ecdc.europa.eu/en/antimicrobial-resistance/surveillance-and-disease-data/data-ecdc) **[7–9]**. ESBL-producing *Enterobacteriaceae* are endemic in different Portuguese clinical settings, and the widespread of main ESBL (CTX-M-15, SHV-12), and subsequently KPC-3, had been linked to the spread of particular *K. pneumoniae* (ST15, ST147, ST348) or *E. coli* clones (ST131) and plasmid types (IncR, IncF, IncHI2) **[7, 9, 10]**.

Our previous survey on qAmpC-producing *Enterobacteriaceae* dates back to the period of 2002 to 2008, and indicated a high occurrence of DHA-1-producing *K. pneumoniae* isolates **[8]**, but detailed molecular analysis was not performed and more recent epidemiological trends are not known. We aim to provide detailed molecular data on clinical qAmpC producing *Enterobacteriaceae* from two different periods (2002-2008 and 2010-2013) in order to clarify the contribution of clonal and plasmid genetic platforms for the current epidemiological scenario concerning extended-spectrum beta-lactams resistance.

## 2. Material and methods

### 2.1. Bacterial isolates

One thousand two hundred and forty-six *Enterobacteriaceae* isolates non-susceptible to third-generation cephalosporins causing community- or hospital-acquired human infections, and belonging to species lacking inducible chromosomal AmpC (icr-AmpC) genes (723 *E. coli*, 486 *K. pneumoniae*, 28 *Klebsiella oxytoca*, 9 *Proteus mirabilis*) were investigated. They were recovered during a 4-year period (2010-2013) from general hospitals located at the North (HA; 63.2%; 2010-2012) and Centre (HD; 5.5%; June 2012-August 2013) of Portugal, and a community laboratory at the North (CL2; 31.4%; August 2012-August 2013), all serving a population of almost 3 000 000 people. Bacterial identification and antibiotic susceptibility testing were performed as described **[8]**.

Additionally, 50 clinical qAmpC producers (38 *K. pneumoniae*, 4 *K. oxytoca*, 3 *Klebsiella variicola*, 5 *E. coli*) identified in our previous survey (2002-2008) **[8]**, at the same clinical institutions, were included for further characterization (clonal identification by multilocus sequence typing and characterization of *bla*_qAmpC_plasmids and surrounding regions) in order to understand the clonal and plasmid dynamics throughout time, and putatively identify persistent genetic entities.

### 2.2. Detection and characterization of isolates carrying qAmpC genes

Presumptive qAmpC producers were preliminarily identified using phenotypic criteria and *bla*_qAmpC_genes were further identified by PCR and sequencing **[8]**. Isolates carrying *bla* _qAmpC_ were also analysed for the presence of ESBL, carbapenemases or plasmid-mediated quinolone resistance genes [PMQR; *qnr* and *aac(6’)-Ib-cr*] **[2, 8, 11]**. *K. variicola*, *E. coli* phylogroups, and *E. coli* O25b-ST131 were identified as described **[10, 12]**. Clonal relatedness was established by *Xba*I-PFGE **[11]**, and MLST for representative *K. pneumoniae* (different PFGE-patterns) (http://bigsdb.pasteur.fr/klebsiella/primers_used.html) and B2- and D-*E. coli* (http://mlst.warwick.ac.uk/mlst/dbs/Ecoli/documents/primersColi_html) isolates. Amplification and sequencing of *wzi* gene was conducted in representative ST11-*K. pneumoniae* (different PFGE-patterns) isolates in order to predict capsular locus (KL) **[13]**.

The location of *bla*_qAmpC_was assessed by hybridization of S1- or I-*Ceu*I-digested genomic DNA **[11]**. Plasmids carrying *bla* _qAmpC_were characterized by replicon typing (PCR, sequencing and hybridization) and subtyping (http://pubmlst.org/plasmid/primers/incI1.shtml) **[2, 11]**. The *bla*_DHA_genetic environment (ca. 22 kb) was investigated by PCR mapping in four representative isolates from different species (1 *E. coli*, 1 *K. oxytoca*, and 1 *K. pneumoniae* producing DHA-1; 1 *K. pneumoniae* producing DHA-6), according to the genetic context previously described in pKPS30 (NC_023314) (Table S1). Additionally, the presence of antibiotic resistance genes within the variable region 1 (VR1) [*catB3*, *aac-(6’)-Ib-cr*, *bla*_OXA-1_, *arr3*] was confirmed by PCR and sequencing in all isolates **[14]**. The genetic surrounding of *bla*_CMY-2_ was characterized by PCR based on previously described platforms **[4]**.

### 2.3. Whole genome sequencing and bioinformatics analysis

WGS of two *K. pneumoniae* isolates of the epidemic ST11-KL105 clone identified in 2006 (strain H642) and 2011 (H1523) was performed by Hi Seq 2000 Sequencing System (Illumina Inc., San Diego, CA, USA) (2 × 125 bp paired-ended reads, coverage 100×). Raw sequence reads were *de novo* assembled using SPAdes version 3.9.0 (http://cab.spbu.ru/software/spades/) and contigs were annotated with Prokka (http://vicbioinformatics.com/). This Whole Genome Shotgun project has been deposited at DDBJ/ENA/GenBank under the accessions QTTC00000000 and QTTD00000000. Ten ST11-KL105 *K. pneumonia* genomes and thirty DHA-1-producing *K. pneumoniae* genomes available on GenBank database (https://www.ncbi.nlm.nih.gov/genbank/) as of July 2018 were included for comparison (Supplementary Table S2 and Table S3).

Identification of acquired antibiotic resistance genes and plasmid replicons was evaluated with ResFinder 3.0 and PlasmidFinder 1.3 tools, respectively (https://cge.cbs.dtu.dk/services/resfinder/, https://cge.cbs.dtu.dk/services/PlasmidFinder/), and KL were identified using Kaptive (https://github.com/katholt/Kaptive). Contigs were compared with known DHA-1-encoding plasmids by progressiveMauve **[15]**. The BLAST Ring Image Generator (BRIG) was used to compare DHA-1 encoding IncR plasmids identified in this study with those deposited on public databases **[16]**.

### 2.4. Statistical analysis

Statistical significance for comparison of proportions was calculated by the χ ^2^test or Fisher exact test using IBM SPSS Statistics 22.0 software (*P* values of <0.05 were considered statistically significant).

## 3. Results and discussion

### 3.1. Occurrence and temporal trends of qAmpC-producing Enterobacteriaceae

The *bla*_qAmpC_genes were detected in 2.6% of the isolates (n=32/1246), corresponding to 16 *E. coli*, 13 *K. pneumoniae*, 2 *K. oxytoca* and 1 *P. mirabilis*, in most cases obtained from urine (44%) or sputum (20%) samples (Table 1). The remaining presumptive qAmpC producers were possible ESBL producers, hyperproducers of chromosomal AmpC enzymes (*E. coli*) or hyperproducers of class A β-lactamases (*Klebsiella* spp.) combined with alterations in porin channels.

**Table 1.**
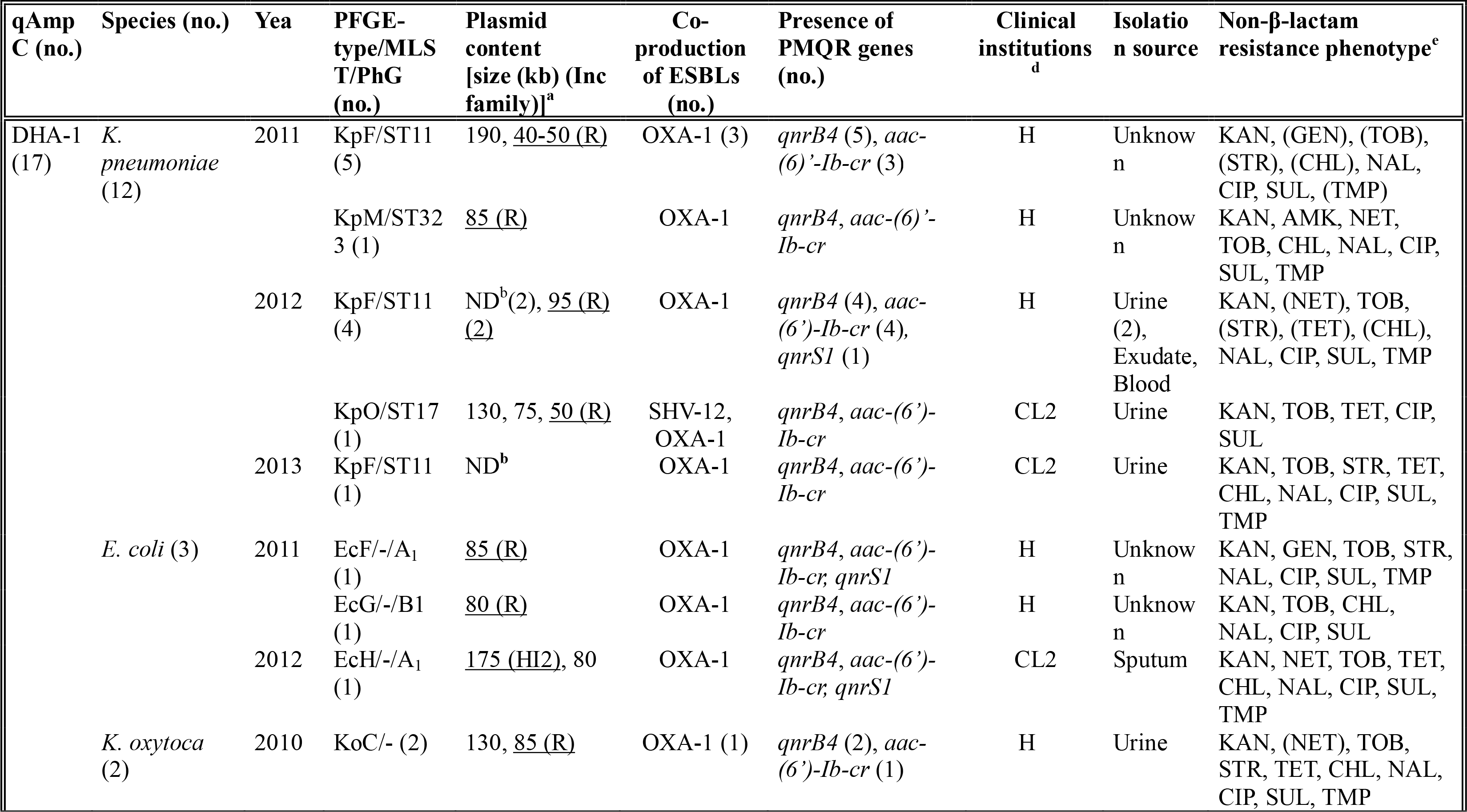
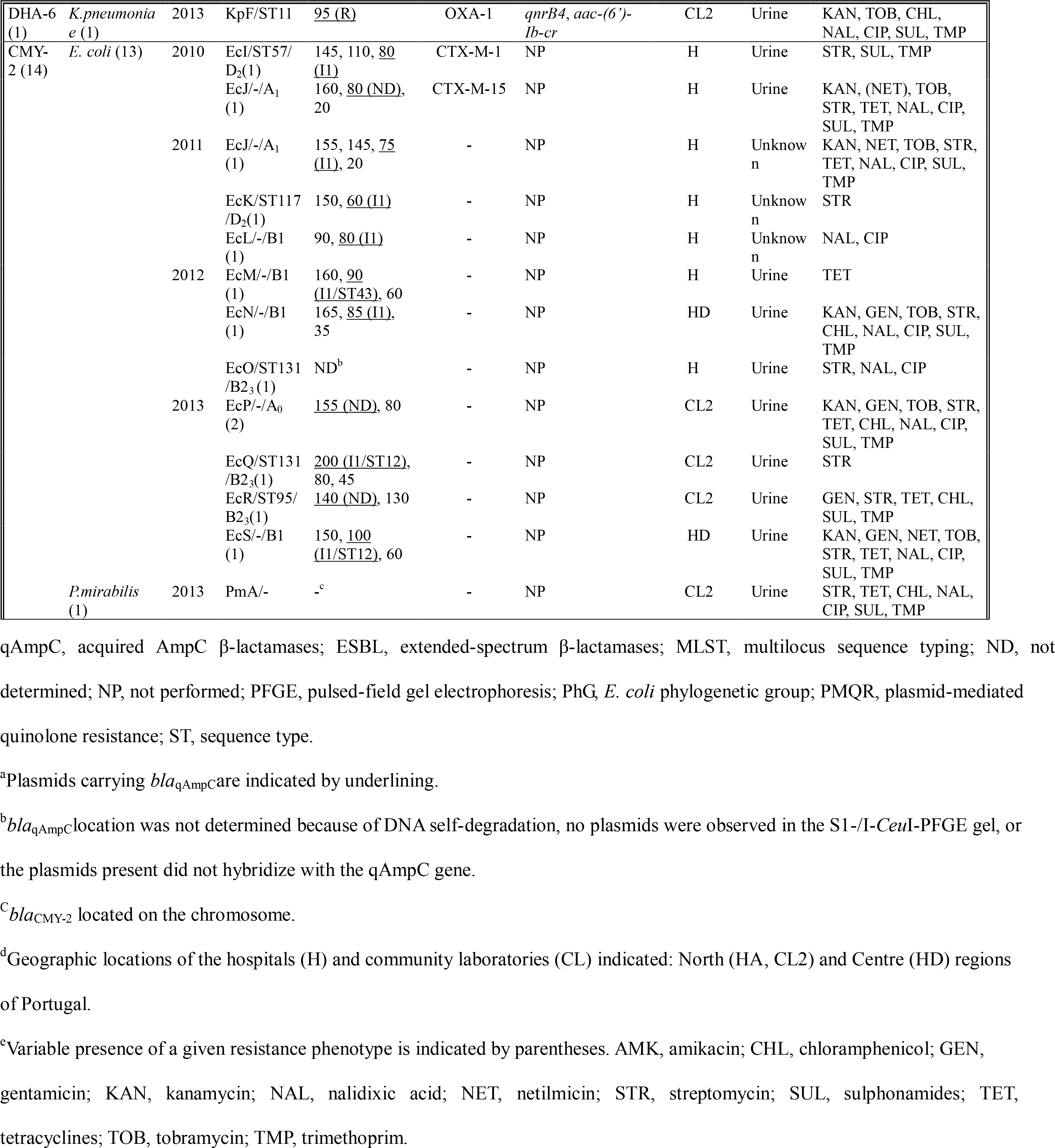
Epidemiological data of qAmpC-producing *Enterobacteriaceae* recovered from Portuguese clinical settings (2010-2013).

The occurrence of qAmpC-producing *Enterobacteriaceae* was lower than that reported previously in a smaller sample (7.4%; 50/675 in 2002-2008) from the same clinical institutions **[8]**, but it is in accordance with rates reported by some European countries (0.06%-1.3% in Spain; 5% in The Netherlands) **[5, 17, 18]**. However, it is of note that most studies are biased towards detection of specific enzymes (most frequently CMY-2 or DHA-1) or species (*E. coli* or *K. pneumoniae*) **[2, 3, 6]**, hindering to establish the overall occurrence of qAmpC producers. Moreover, our study revealed similar occurrence rates of qAmpC-producing *E. coli* (2.2%, n=16/723) or *K. pneumoniae* (2.7%, n=13/486) to those described in Spain, Canada, or Iran (0.8%-3.6%) **[2, 3, 20]**. Nevertheless, qAmpC occurrence was much lower than that of ESBL-producing *Enterobacteriaceae* reported in our country for the same time period (13.4-44.4% of ESBL-producing *E. coli*; 39.0-82.5% of ESBL-producing *K. pneumoniae*) **[20]**. The proportion of qAmpC producers slightly varied over time in hospitals (2.5% in 2010, 3.6% in 2011, 2.0% in 2012, and 2.5% in 2013) or community (1.5% in 2012, and 2.7% in 2013), although the number and type of institutions analysed varied during the whole period.

In this study, qAmpC enzymes identified were the same than those circulating previously (DHA-1 and CMY-2) though in different proportions, with exception of DHA-6 reported in Portugal for the first time. DHA-1 represented 53% (n=17/32) of the qAmpCs identified between 2010 and 2013, a lower frequency than that observed in 2002-2008 (94%; *p*<0.000001) **[8]**, the latter strongly biased by the occurrence of *K. pneumoniae* hospital outbreaks. It was also identified in *E. coli* and *K. oxytoca* (Table 1), highlighting the continuous dissemination of DHA-1 in species other than *K. pneumoniae*, as previously observed **[8]**. On the contrary, CMY-2 rates increased (44%, n=14/32) compared with those reported in 2002-2008 (6%; *p*<0.000001) **[8]**, associated in all but one case with *E. coli* mostly recovered from hospitalized patients (61.5%) (Table 1). DHA-6 (3%, n=1/32; differing from DHA-1 in A226T mutation) was identified in one *K. pneumoniae* (2013; CL2). Interestingly, the persistence of DHA-1 through time and the recent increase of CMY-2 in clinical institutions differ from the worldwide scenario reporting CMY-2 as the dominant qAmpC among clinical *Enterobacteriaceae* **[1, 4, 21]**.

### 3.2. Co-carriage of qAmpC and other antibiotic resistance genes

The majority of DHA-1 producers co-expressed OXA-1 (82%), while CMY-2 producers co-expressed occasionally CTX-M-1 or CTX-M-15 (14%) (Table 1). Carbapenemase genes were not detected, despite some DHA-producers (8 *K. pneumoniae*, 1 *E. coli*) were resistant to ertapenem and one CMY-2-producing *E. coli* was resistant to imipenem, which suggests the involvement of permeability defects **[2, 22]**. DHA-1-producers also harboured frequently PMQR genes [*qnrB4*, 100%; *aac-(6’)-Ib-cr*, 82%; *qnrS1*, 18%], being the high occurrence of *qnrB4* and *aac-(6’)-Ib-cr* explained by their genetic vicinity to the *bla*_DHA-1_ (see section 3.4.) **[1, 2]**. The frequent detection of isolates carrying qAmpC and PMQR, and exhibiting a multidrug-resistant (MDR) phenotype greatly limits effective therapeutic options in infections caused by qAmpC-producing *Enterobacteriaceae*.

### 3.3. Clonal relatedness among qAmpC-producing Enterobacteriaceae

During the first period analysed (2002-2008), *K. pneumoniae* isolates were associated with a high diversity of clones (11 PFGE types) belonging to 10 STs (ST11, ST440, ST416, ST443, ST1871, ST1872, ST1873, ST1874, ST1224, ST1380) (Table 2), most of them observed sporadically. Two MDR *K. pneumoniae* clones were frequently identified: (i) ST1380 (37%; clone KpB), associated with a small outbreak in 2004, producing DHA-1 and diverse SHV-ESBLs; (ii) ST11 (34%; clone KpF), harbouring *wzi*75 and the capsular type KL105, detected in different hospitals between 2006 and 2008, producing DHA-1 and OXA-1 (Table 2). Remarkably, three DHA-1 producing isolates reported in 2007 in different wards (paediatrics, medicine) from the same hospital were re-identified as *K variicola* (Table 2), a species increasingly recognized as a frequent cause of human clinical infections that can be misidentified by automated methods, including MALDI-TOF MS **[23]**. More recently (2010-2013), the population of *K. pneumoniae* producing DHA was dominated by the MDR *K. pneumoniae* ST11-KL105 (KpF clone) (n=11; 85%), that persisted till the end of the period analysed at least in Hospital A (Table 1). The identification of this lineage causing community acquired infections (this study) or colonizing the gastrointestinal tract of long-term care facilities residents, suggests a wider and current dissemination **[24]**.

**Table 2.**
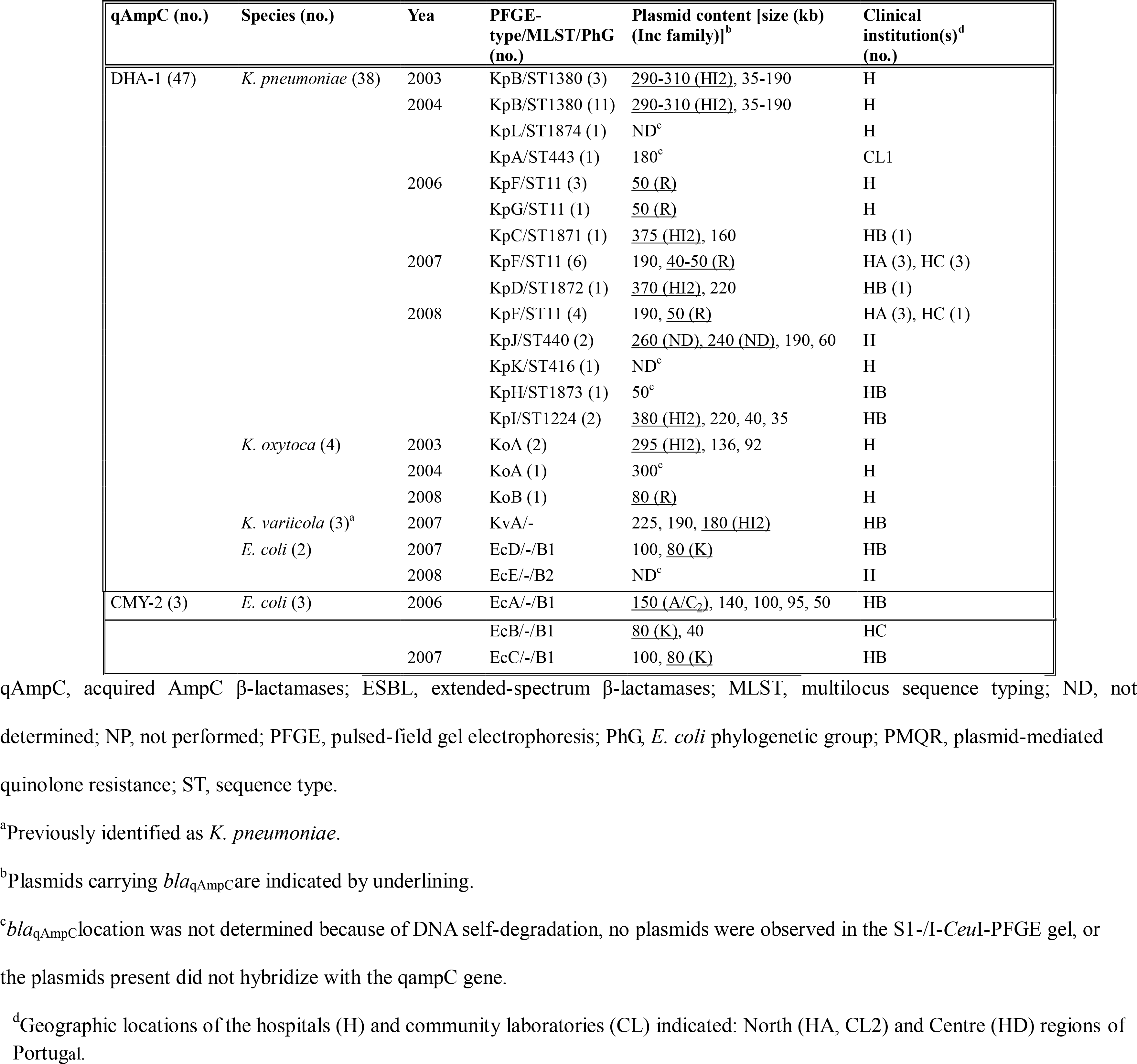
Characterisation of plasmid and population structure of qAmpC-producing *Enterobacteriaceae* previously recovered from Portuguese clinical settings (2002-2008) [8].

| qAmpC (no.) | Species (no.) | Yea | PFGE-type/MLST/PhG<br>(no.) | Plasmid content [size (kb)<br>(Inc family)]<sup>b</sup> | Clinical<br>institution(s)<sup>d</sup><br>(no.) |
| --- | --- | --- | --- | --- | --- |
| DHA-1 (47) | *K. pneumoniae* (38) | 2003 | KpB/ST1380 (3) | <u>290-310 (HI2)</u>, 35-190 | H |
|  |  | 2004 | KpB/ST1380 (11) | <u>290-310 (HI2)</u>, 35-190 | H |
|  |  |  | KpL/ST1874 (1) | ND<sup>c</sup> | H |
|  |  |  | KpA/ST443 (1) | 180<sup>c</sup> | CL1 |
|  |  | 2006 | KpF/ST11 (3) | <u>50 (R)</u> | H |
|  |  |  | KpG/ST11 (1) | <u>50 (R)</u> | H |
|  |  |  | KpC/ST1871 (1) | <u>375 (HI2)</u>, 160 | HB (1) |
|  |  | 2007 | KpF/ST11 (6) | 190, <u>40-50 (R)</u> | HA (3), HC (3) |
|  |  |  | KpD/ST1872 (1) | <u>370 (HI2)</u>, 220 | HB (1) |
|  |  | 2008 | KpF/ST11 (4) | 190, <u>50 (R)</u> | HA (3), HC (1) |
|  |  |  | KpJ/ST440 (2) | <u>260 (ND)</u>, <u>240 (ND)</u>, 190, 60 | H |
|  |  |  | KpK/ST416 (1) | ND<sup>c</sup> | H |
|  |  |  | KpH/ST1873 (1) | 50<sup>c</sup> | HB |
|  |  |  | KpI/ST1224 (2) | <u>380 (HI2)</u>, 220, 40, 35 | HB |
|  | *K. oxytoca* (4) | 2003 | KoA (2) | <u>295 (HI2)</u>, 136, 92 | H |
|  |  | 2004 | KoA (1) | 300<sup>c</sup> | H |
|  |  | 2008 | KoB (1) | <u>80 (R)</u> | H |
|  | *K. variicola* (3)<sup>a</sup> | 2007 | KvA/- | 225, 190, <u>180 (HI2)</u> | HB |
|  | *E. coli* (2) | 2007 | EcD/-/B1 | 100, <u>80 (K)</u> | HB |
|  |  | 2008 | EcE/-/B2 | ND<sup>c</sup> | H |
| CMY-2 (3) | *E. coli* (3) | 2006 | EcA/-/B1 | <u>150 (A/C<sub>2</sub>)</u>, 140, 100, 95, 50 | HB |
|  |  |  | EcB/-/B1 | <u>80 (K)</u>, 40 | HC |
|  |  | 2007 | EcC/-/B1 | 100, <u>80 (K)</u> | HB |

qAmpC, acquired AmpC β-lactamases; ESBL, extended-spectrum β-lactamases; MLST, multilocus sequence typing; ND, not determined; NP, not performed; PFGE, pulsed-field gel electrophoresis; PhG, *E. coli* phylogenetic group; PMQR, plasmid-mediated quinolone resistance; ST, sequence type.
<sup>a</sup>Previously identified as *K. pneumoniae*.
<sup>b</sup>Plasmids carrying *bla*<sub>qAmpC</sub> are indicated by underlining.
<sup>c</sup>*bla*<sub>qAmpC</sub> location was not determined because of DNA self-degradation, no plasmids were observed in the S1-/I-CeuI-PFGE gel, or the plasmids present did not hybridize with the qampC gene.
<sup>d</sup>Geographic locations of the hospitals (H) and community laboratories (CL) indicated: North (HA, CL2) and Centre (HD) regions of Portugal.

Notably, this ST11-KL105 *K. pneumoniae* lineage was not detected among ESBL- or carbapenemase-producers circulating in the same clinical institutions and period of time **[7, 9]**, indicating that the selection of ESBL or qAmpC producing *K. pneumoniae* occurred independently in different genetic backgrounds, or an eventual importation event of a DHA-1-producing ST11 *K. pneumoniae* strain. In fact, ST11-KL105 isolates producing DHA-1 have been reported in different countries, either causing infection or linked to human colonization (Table S2). These isolates carried often other antibiotic resistance genes [*aac(6’)-Ib-cr*, *qnrB4*, *aph(3’)-Ia*, *catA1*, *catB*, *arr-3*, *aadA2*, *dfrA*] and plasmid replicons (mostly FII_K_and R), identified also in our isolates (data not shown) (Table S3). Furthermore, ST11-KL105 strains carrying other determinants of β-lactam resistance (*bla*_CTX-M-15_or *bla* _KPC-2_) have also been identified in different countries.

One additional ST11-*K. pneumoniae* isolate with *wzi*24 presumptively associated with K24 was identified in 2006 (PFGE-type KpG) (Table 2). One could hypothesize that it could have emerged by recombination events involving also the capsular locus or it could eventually correspond to a different ST11 lineage. In fact, different ST11 lineages carrying specific KL types have been identified suggesting a suboptimal resolution of MLST **[25]**. Other sporadic *K. pneumoniae* clones identified in this study were ST17 (belonging to clonal group 17) and ST323 (Table 1). The genomes of DHA-1-producing strains deposited in public databases, most of them from 2010 onwards, belong to a high diversity of clones and capsular types suggesting a high diversity of genetic backgrounds circulating worldwide (Table S3).

A high diversity of CMY-2-producing *E. coli* isolates belonging to phylogroups A (n=4; 31%), B1 (n=4; 31%), B2_3_ (n=3; 23%) or D (n=2; 15%) (Table 1) was observed, probably reflecting independent acquisition events, as reported for ESBL producers **[10]**. The detection of CMY-2 amongst the widespread B2-ST131 (n=2), B2-ST95 (n=1), D-ST57 (n=1) and D-ST117 (n=1) clones, frequently detected as ESBL producers in the same clinical institutions **[10]**, corroborates the ability of these clones to acquire a diversity of antimicrobial resistance platforms.

### 3.4. Location of bla_qAmpC_ on specific plasmid types in different species

The characterization of 82 strains revealed that qAmpC genes (64 *bla*_DHA-1_, 17 *bla*_CMY-2_, 1 *bla*_DHA-6_) were located on plasmids of various sizes (n=71/82, 86.6%, 40-380 kb) or on the chromosome (n=1/82, 1.2%), while in 12.2% (n=10/82) of the isolates its specific location was not determined (Table 1 and Table 2).

Plasmids carrying *bla*_DHA-1_ were identified as 40-95 kb IncR (n=28) or 290-380 kb IncHI2 (n=24) in different *Enterobacteriaceae* species, 80 kb IncK (n=1) in *E. coli* and 240-260 kb non-typeable (n=2) plasmids in *K. pneumoniae*, corroborating previous data **[26]**. It is noteworthy the shift observed in plasmid types carrying *bla*_DHA-1_over time (Table 1 and Table 2). IncHI2 were dominant between 2002 and 2008 (n=21) and circulated in different *K. pneumoniae* clones, *K. oxytoca* and *K. variicola,* but were rarely identified in the recent period (2010-2013). On the other hand, IncR plasmids were identified since 2006 in a ST11-*K. pneumoniae* clone, and persisted not only in this clone but also in other MDR *K. pneumoniae* clones (ST17 and ST323), *E. coli* or *K. oxytoca* (Table 1).

IncR plasmids have been found in different epidemic *K. pneumoniae* clones in Portugal associated with distinct acquired β-lactamases (e.g. ST15, ST147 and ST336 carrying CTX-M-15 or SHV-12) in the same clinical institution (HA) included in this study **[7]**, further highlighting a well adaptation to this species **[27, 28]**. Besides, IncR plasmids have also been associated with carriage of genes encoding carbapenemases (VIM-1, KPC-2, NDM-1), aminoglycosides (*armA*, *rmtB*) or fluoroquinolones [*qnrB4/6*, *qnrS1*, *aac(6’)-Ib-cr*] resistance **[7, 26, 27]**.

Comparison of IncR plasmids identified in this study (pH642 and pH1523) with other representative IncR plasmids, namely: pKPS30 (NC_023314), pCNR3 (LT994834), and pCNR95 (LT994839) isolated from ST11-*K. pneumoniae* strains in France, and pYDC676 (KT225462) isolated from a ST37-*K. pneumoniae* in the USA showed a high identity of plasmid backbones (*repB*, *resD*, *parAB*, *umuCD*, *retA*, *vagDC* genes) (Figure 1) **[27]**. Moreover, the ca. 22 kb multidrug resistance region (MRR), comprising the *bla*_DHA-1_ gene, was highly similar in all plasmids except pYDC676 (that seems to have lost a part of it), and identical to that previously described in pKPS30 (NC_023314), the first complete plasmid-mediated DHA-1 beta-lactamase belonging to IncR group in a *K. pneumoniae* in 2008 **[26]**.

**Figure 1.**
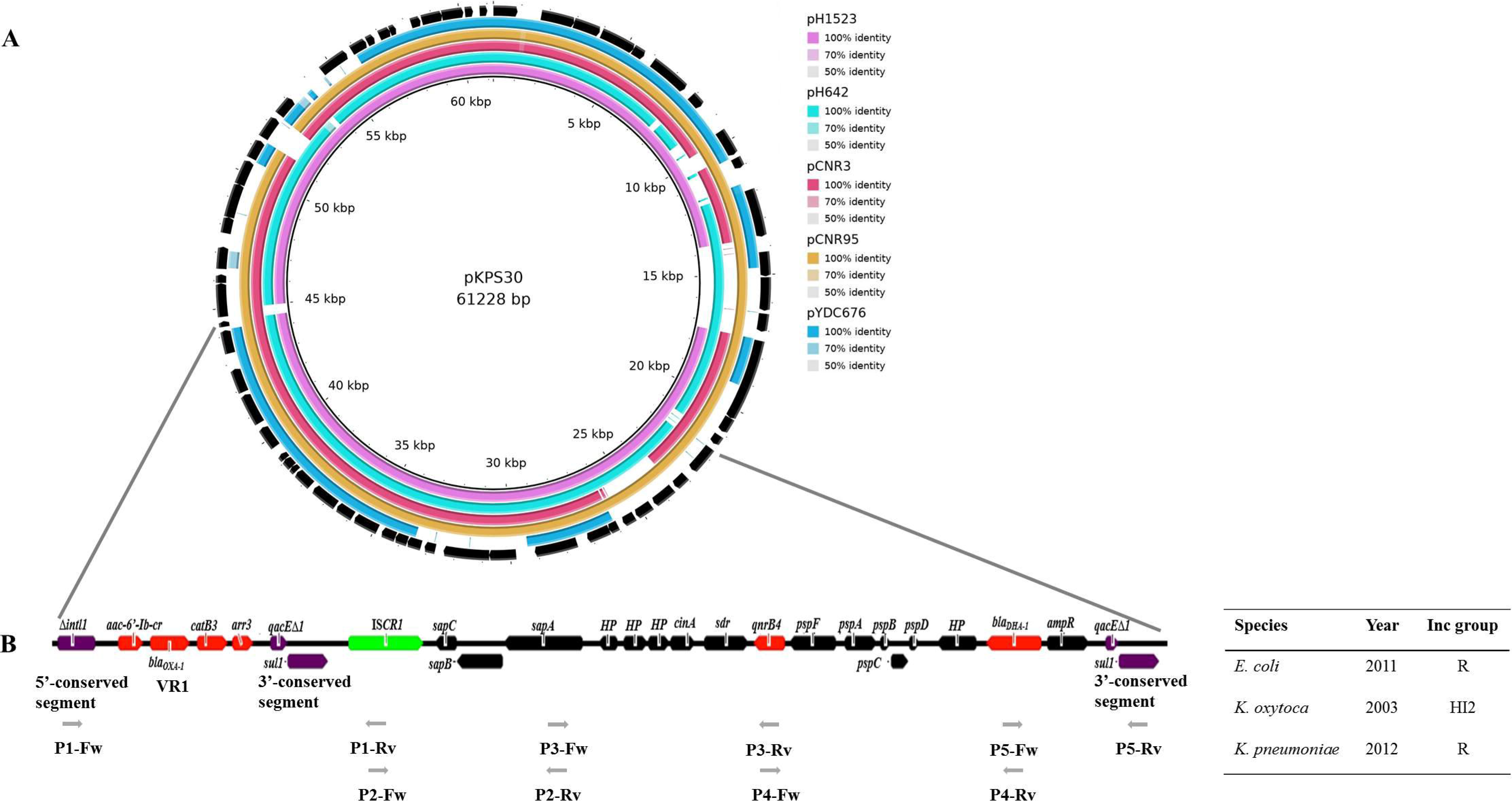
A) Comparative analysis of representative DHA-1–harbouring IncR plasmids from different human isolation sources and geographic regions by BRIG. pKPS30 (ST11-*K. pneumoniae*, NC_023314) was used as a reference plasmid, GenBank accession numbers of pYDC676 (ST37-*K. pneumoniae*), pCNR3 (ST11-*K. pneumoniae*), and pCNR95 (ST11-*K. pneumoniae*) are KT225462, LT994834, and LT994839, respectively. The outermost circle shows the direction of transcriptional open-reading frames of the reference plasmid. **B) Genetic context of *bla*_DHA-1_.**The *int*, encoding the integrase (5’-conserved segment) and *qacEΔ1sul1* (3’-conserved segment) genes (purple), other genes (black), antimicrobial drug resistance genes (red), and insertion sequence (green) are indicated. The strategy for PCR mapping of *bla*_DHA-1_genetic environment (ca. 22 kb) is indicated by grey arrows. Primer (P) sequences are listed in Supplementary data (Table S1).

This region includes an unusual class 1 integron of *sul1*-type which contains two partial copies of the 3’-conserved segment (3’CS) *qacEΔsul1* surrounding a common IS*CR1* region and *qnrB4*, *sap* (encoding ABC transporter family) and *psp* (encoding phage shock proteins) operons (Figure 1B) **[26]**. A similar genetic environment was observed for *bla* _DHA-6_(data not shown), that was located at a 50 kb IncR plasmid in one ST11-KL105 *K. pneumoniae* strain, suggesting *in vivo* evolution. This MRR was also observed in plasmids belonging to different Inc groups (IncR and IncHI2) and in different *Enterobacteriaceae* species from our collection (data not shown), with slight differences in the integron variable region. Interestingly, all but one DHA-1-producing *Enterobacteriaceae* collected before 2005 (n=18/19) contained only *aac(6′)-Ib-cr* in IncHI2 plasmids, whereas most DHA-1 producers identified afterwards (n=44/47) carried a larger gene cassette [*aac(6′)-Ib-cr-bla*_OXA-1_-*catB3*-*arr3*] in either IncHI2 or IncR plasmids.

On the other hand, the few *E. coli* isolates from phylogenetic group B1 identified in 2006 and 2007 carried *bla*_CMY-2_ in IncK or IncA/C _2_plasmids, that are considered main vehicles of this gene besides IncI1 **[4]**. More recently (2010-2013), *bla*_CMY-2_ was mostly linked to IncI1 plasmids in diverse *E. coli* isolates (62%) causing hospital or community acquired infections, being in agreement with reports from Spain **[1, 4]**, or in the chromosome of *P. mirabilis* (n=1) (Table 1). Two different IncI1 subtypes were identified, namely the widely disseminated IncI1/ST12 (60-200 kb; n=2; HD and CL2) (https://pubmlst.org/bigsdb?db=pubmlst_plasmid_isolates&page=profiles) and IncI1/ST43 (90 kb; n=1; HA) (Table 1), the latter previously identified in two CMY-2-producing *E. coli* isolates from the United Kingdom (https://pubmlst.org/bigsdb?db=pubmlst_plasmid_isolates&page=profiles). It is of note that the IncI1/ST12 subtype carrying *bla*_CMY-2_ has been frequently detected in *E. coli* or *Salmonella* sp. isolated from food animals (mainly poultry) or meat from different countries, including in Portugal (https://pubmlst.org/bigsdb?db=pubmlst_plasmid_isolates&page=profiles) **[29, 30]**, suggesting transmission to humans through the food chain.

The *bla*_CMY-2_ was in nearly all *E. coli* isolates linked to the transposon-like element IS*Ecp1* (n=12/17; IS*Ecp1*/ΔIS*Ecp1*-*bla*_CMY-2_-*blc*-*sugE*), which corresponds to the genetic context most frequently described **[1, 4]**. The *int* gene from the SXT/R391 integrative and conjugative element (ICE) family was identified in the *P. mirabilis* isolate harbouring a chromosomally-encoded CMY-2, supporting the involvement of ICEs in the mobilization of *bla* _CMY-2_ (data not shown) **[31]**.

## 4. Conclusions

This work constitutes one of the most comprehensive and detailed studies analysing the molecular epidemiology of qAmpC among clinical *Enterobacteriaceae*. We highlight significant epidemiological changes throughout time on bacterial hosts and plasmid drivers of *bla*_CMY-2_ and *bla* _DHA-1_, towards the selection of a specific *K. pneumoniae* lineage carrying DHA-1 (ST11-KL105) and plasmids worldwide disseminated (IncR-DHA-1; IncI1/ST12-CMY-2). Finally, we hypothesized that genetic backgrounds (clones, plasmids, integron platforms) differing from those responsible for ESBL spread in the same institutions and time periods in our country, provided bacterial features that might have contributed to the differential expansion of ESBL and qAmpC in Portugal.

## Acknowledgments

We are grateful to the healthcare and management personnel of the hospitals and clinical laboratories enrolled in this study, and to João A. Carriço for the support in bioinformatics tools. TGR performed the experiments, analysis and interpretation of data, and wrote the manuscript. ÂN orientated the study, contributed to the analysis and interpretation of data, and critically reviewed the manuscript. CR contributed to WGS and statistical analysis. RN and FF performed part of the experiments. EM and LP conceived the study and revised it critically.

## Declarations

### Funding

This work was supported by European Union (FEDER funds) through Programa Operacional Fatores de Competitividade – COMPETE, and by National Funds [Fundação para a Ciência e a Tecnologia (FCT)] [POCI/01/0145/FEDER/007728 and UID/Multi/04378/2013]. T. G. R., Â. N. and C. R. were supported by FCT fellowships [SFRH/BD/75752/2011, SFRH/BPD/104927/2014 and SFRH/BD/84341/2012, respectively] through Programa Operacional Capital Humano (POPH).

### Competing Interests

The authors declare no conflicts of interest.

### Ethical Approval

Not required

## Abbreviations

CS: conserved segment
ESBL: extended-spectrum β-lactamases
ICE: integrative and conjugative element
icr-AmpC: inducible chromosomal AmpC
KL: capsular locus
MDR: multidrug-resistant
MRR: multidrug resistance region
PMQR: plasmid-mediated quinolone resistance
qAmpC: acquired AmpC β-lactamases
VR1: variable region 1.

