## Supplementary data for "Significant temporal shifts on clonal and plasmid backgrounds of *Enterobacteriaceae* producing acquired AmpC in Portuguese clinical settings"

**TABLE S1. Sequences of oligonucleotides used for PCR mapping.**

| **Primer** | **Name** | **Oligonucleotide sequence (5’ to 3’)** | **Gene or target** | **Expected Amplicon size (bp)** | **References** |
| --- | --- | --- | --- | --- | --- |
| P1-Fw | Intl1-R | TTCGAATGTCGTAACCGC | *intl1* | 6490 | 1 |
| P1-Rv | orf513-R | CTGAGGGTGTGAGCGAG | IS*CR1* |  |  |
| P2-Fw | orf513-F | ATGGTTTCATGCGGGTT | *ISCR1* | 3798 | 1 |
| P2-Rv | sapA-F | CAGTGGGTTCGTCTATTGC | *sapA* |  |  |
| P3-Fw | sapA3-R | GCAGCAGCCTGACCACTC | *sapA* | 4872 | 1 |
| P3-Rv | qnrB-F | CCGACCTGAGCGGCACTGA | *qnrB* |  |  |
| P4-Fw | qnrB-R | CGCTCCATGAGCAACGATGCCT | *qnrB* | 5782 | 1 |
| P4-Rv | AmpC/III_Fw | CATTAAACCGCTGATGGCAC | *bla*_DHA_ |  |  |
| P5-Fw | AmpC/III_Rev | GCTTTGACTCTTTCGGTATTCG | *bla*_DHA_ | 2925 | 2 |
| P5-Rv | sul5′ | ATCAGATGCACCGTGTTTCA | *sul1* |  | 3 |

**Table S2. ST11-*Klebsiella pneumoniae* KL105 genomes publicly available on GenBank database (**[**www.ncbi.nlm.nih.gov/genbank/**](http://www.ncbi.nlm.nih.gov/genbank/)**).**

| **Strain** | **Year** | **Country** | | **Genes coding for resistance to different antibiotic classes (ResFinder)** | | | | | | | **Incompatibility group (s)**  **(Plasmid Finder)^a^** | | **Infection (I) or Colonization (C)** | | **GenBank Accessions** |
| --- | --- | --- | --- | --- | --- | --- | --- | --- | --- | --- | --- | --- | --- | --- | --- |
|  |  |  |  | **Beta-lactams** | **Aminoglycosides** | | | **Quinolo**  **nes** | | **Others (nº.)** |  |  |  |  |  |
| DU4033/04 | 2004 | Singapore | *bla*_DHA-1_, *bla*_SHV-11_, *bla*_OXA-1_ | | | *aac(6')-Ib-cr*, *aadA2*, *aph(3')-Ia* | *qnrB4*, *oqxAB* | | *fosA, mph(A), catA1, catB4,arr-3, sul1, tet(A), dfrA12* | | | FIB(K), R, FII(K), ColRNAI | I | ERX2005511 | |
| DU10252/04 | 2004 | Singapore | *bla*_TEM-1B_, *bla*_SHV-12_, *bla*_OXA-1_ | | | *aac(6')-Ib-cr*, *aadA2*, *aph(3')-Ia* | *oqxAB* | | *fosA, mph(A), catA1, catB4,arr-3, sul1, dfrA12* | | | FIB(K), R, X3 | I | ERX2005514 | |
| DU38032/05 | 2005 | Singapore | *bla*_CTX-M-15_, *bla*_SHV-11_, *bla*_OXA-1_ | | | *aac(3)-IIa*, *aac(6')-Ib-cr*, *aadA2*, *aph(3')-Ia* | *qnrA1*, *oqxAB* | | *fosA* | | | FIB(K), R, FII(K), ColRNAI, FII | I | ERX2005518 | |
| H642 | 2006 | Portugal | *bla*_DHA-1_, *bla*_SHV-11_, *bla*_OXA-1_ | | | *aac(6')-Ib-cr, aadA2, aph(3')-Ia* | *qnrB4, oqxAB* | | *fosA*, *mph(A)*, *catA*1, *catB3*, *arr-3*, Δ*sul1* (2), *tet(A)*, *drfA12* | | | FIB(K), R, FII(K) | I | QTTC00000000 | |
| H1523 | 2011 | Portugal | *bla*_DHA-1,_ *bla*_TEM-1B_, *bla*_SHV-11_, *bla*_OXA-1_ | | | *aac(6')-Ib-cr, aadA2, aph(3')-Ia, aph(4)-Ib* | *qnrB4, oqxAB* | | *fosA*, *mph(A)*, *catA1*, *catB3*, *arr-3*, Δ*sul1* (2), *tet(A)*, *drfA25* | | | FIB(K), R, FII(K), N | I | QTTD00000000 | |
| IS33 | 2011 | Austria | *bla*_DHA-1_, *bla*_CTX-M-15_, *bla*_TEM-1A_, *bla*_SHV-11_, *bla*_OXA-9_ | | | *aac(3)-IIa*, *aac(6')-Ib-cr*, *aadA1*, *aadA2* | *qnrB4*, *oqxAB* | | *fosA*, *catA1*, Δ*sul1* (2) | | | FIB(K), R, FII(K) | I | CBWI00000000.1 | |
| AUH-KIMP217 | 2013 | Lebanon | *bla*_DHA-1_, *bla*_NDM-1_, *bla*_CTX-M-15_, *bla*_TEM-1B_, *bla*_SHV-11_, *bla*_OXA-1/-9_ | | | *aac(6')-Ib*, *aadA1*, *aadA2*, *aph(3')-Ia*, *aph(3')-VI* | *qnrB4*, *qnrS1*, *oqxAB* | | fosA, mph(A), catA1, catB4, *arr-3*, sul1 (2), tet(A), dfrA12 | | | FIB(K), R, FII(K), FIB(pQil) | NI | NICN01000000.1 | |
| KPC05 | 2014 | Brazil | *bla*_CTX-M-15_, *bla*_SHV-11_, *bla*_OXA-1_ | | | *aac(6')-Ib-cr, aph(3')-Ia, aph(3')-VIa* | *qnrB19*, *oqxAB* | | *fosA, mph(A),catB4,arr-3, sul1* | | | FIB(K), R, L/M, Q1 | C | NTGJ00000000.1 | |
| KPC40 | 2015 | Brazil | *bla*_CTX-M-15_, *bla*_SHV-11_ | | |  | *oqxAB* | | *fosA* | | | - | I | NIRG00000000.1 | |
| KPC45 | 2015 | Brazil | *bla*_CTX-M-15_, *bla*_SHV-11_ | | |  | *oqxAB* | | *fosA*, *sul1* (2) | | | R | C | LXTP00000000.1 | |
| XL-1 | 2015 | China | *bla*_KPC-2,_ *bla*_SHV-11_ | | | *aac(3)-IVa*, *aadA2*, *aph(4)-Ia* | *oqxAB* | | *fosA*, *sul1*, *drfA12* | | | R, FII(K), ColRNAI | I | MNAK00000000.1 | |
| L37 | 2016 | China | *bla*_KPC-2_, *bla*_SHV-11_ | | | *aadA2* | *oqxAB* | | *fosA*, *sul1* | | | R, ColRNAI | C | NLDT00000000.1 | |

**^a^**Plasmids carrying bla_DHA-1_ are underlined.

**Table S3. DHA-1-producing *K. pneumoniae* genomes publicly available on GenBank database (**[**www.ncbi.nlm.nih.gov/genbank/**](http://www.ncbi.nlm.nih.gov/genbank/)**).**

| **Strain** | **MLST** | **K locus (Kaptive)** | **Year** | **Continent/Country** | **Isolation source** | **GenBank Accessions** |
| --- | --- | --- | --- | --- | --- | --- |
| ARPG-281 | 1 | 19 | 2013 | Asia/Philippines | Blood | NBOH00000000.1 |
| ARPG-315 | 1 | 19 | 2013 | Asia/Philippines | wound abscess | NBOI00000000.1 |
| WCHKP85522 | 1 | 45 | 2015 | Asia/China | NI | NBXV00000000.1 |
| YMC2011/11/B7578 | 11 | 15 | 2011 | Asia/South Korea | Blood | LYPT00000000.1 |
| PB450 | 11 | 15 | 2015 | Asia/Thailand | Urine | FLVC00000000.1 |
| kp10 | 11 | 64 | 2011 | Asia/China | Blood | MNCK00000000.1 |
| AR_0079 | 11 | 64 | NI | America/USA | NI | CP028994.1 |
| TRKP1 K23 | 15 | 19 | 2017 | Asia/China | Urine | NOJX00000000.1 |
| SCKP-LL83 | 15 | 106 | 2017 | Asia/China | NI | NOKM00000000.1 |
| CR48 | 16 | 51 | 2012 | Europe/France | Urine | LS399318.1 |
| PB283 | 29 | 19 | 2015 | Asia/Thailand | Pus | FLVS00000000.1 |
| k2328 | 29 | 30 | 2010 | Europe/UK | Blood | FLGH00000000.1 |
| SCLZ15-011 | 29 | 54 | 2014 | Asia/China | NI | LWLM00000000.1 |
| MRY10-808 | 35 | 62 | 2010 | Asia/Japan | NI | BDLF00000000.1 |
| KPCRETH10 | 37 | 38 | 2015 | Asia/Thailand | Rectal swab | NQAU00000000.1 |
| KpQ24 | 37 | 149 | 2008 | Europe/Spain | Blood | AWON00000000.1 |
| KpQ15 | 37 | 149 | 2008 | Europe/Spain | Blood | AWOM00000000.1 |
| KpQ3 | 37 | 149 | 2008 | Europe/Spain | Blood | AMSU00000000.1 |
| WCHKP030295 | 37 | 158 | 2016 | Asia/China | NI | NWEP00000000.1 |
| SCKP020052 | 54 | 14 | 2016 | Asia/China | NI | PWAS00000000.1 |
| PB209 | 199 | 154 | 2015 | Asia/Thailand | Pus | FLVD00000000.1 |
| PB14 | 234 | 122 | 2015 | Asia/Thailand | Sputum | FLVD00000000.1 |
| 12-3578 | 421 | 123 | 2012 | Asia/China | Blood | AQOC00000000.1 |
| MRY10-848 | 485 | 10 | 2010 | Asia/Japan | NI | BDLG00000000.1 |
| KPPSTH07 | 1307 | 127 | 2016 | Asia/Thailand | Sputum | NQEO00000000.1 |
| SCKP020146 | 2407 | 25 | 2017 | Asia/China | NI | NWFC00000000.1 |
| SCKP040067 | 2407 | 25 | 2017 | Asia/China | NI | PWAN00000000.1 |
| PB517 | Novel ST | 127 | 2015 | Asia/Thailand | Pus | FLXA00000000.1 |
| KPPSTH02 | ND | 9 | 2016 | Asia/Thailand | Sputum | NQEJ00000000.1 |
| 42195 42195 | ND | 14 | 2013 | America/USA | Blood | NDEI00000000.1 |
